# The benefits of insect-swarm hunting to echolocating bats, and its influence on the evolution of bat echolocation signals

**DOI:** 10.1101/554055

**Authors:** Arjan Boonman, Brock Fenton, Yossi Yovel

**Affiliations:** Tel Aviv University; Western University

## Abstract

Predation on swarms of prey, especially using visual information, has drawn much interest in studies of collective movement. Surprisingly, in the field of biosonar this aspect of prey detection, which is probably very common, has received little to no attention. Here, we combine computer simulations and actual echo measurements to accurately estimate the echo intensity of insect swarms of different size and density. We show that swarm echo intensity increases with 3dB for every doubling of insect number, irrespective of swarm density. Thus swarms will be much easier to detect than single insects. Many of the insects bats eat are so small that they are only detectable by echolocation at very short distances. By focusing on detection of swarms of insects, a bat may increase its operating range and diversify its diet. Interestingly, interference between the sound waves reflected from a swarm of insects can sometimes result in echoes that are much weaker than echoes from single insects. We show that bats can reduce this problem by increasing the bandwidth of their echolocation calls. Specifically, a bandwidth of 3-8 kHz would guarantee receiving loud echoes from any angle relative to the swarm. Indeed, many bat species, and specifically bats hunting in open spaces, where swarms are abundant, use echolocation signals with a bandwidth of several kHz. Our results might also explain how the first echolocating bats that probably had limited echolocation abilities, could detect insects through swarm hunting.

## Introduction

Hunting on animal aggregations has been studied from many perspectives including the potential benefits for the prey ([1]) and the challenges faced by the predator ([2]). Surprisingly, one of the groups that most heavily rely on swarm-hunting, namely echolocating bats, have hardly been studied in this context. Previous work on the sensory aspects of swarm hunting concentrated on vision with no research looking at the echoes generated by a swarm ([3], [4]). Previous research on insect echo detection focused on echoes of single insects [5], [6], [7]. In this study, we examine the benefits of hunting for insect swarms using echolocation. Current technological advances enabled us to estimate the reflections produced by even the tiniest mosquito, as well as a swarm of hundreds of insects from any angle. We were thus able to generate natural echo-scenarios that hunting bats will encounter. We use both modeling as well as actual echo measurements to quantify the benefits of hunting for swarms using echolocation. We also analyze the type of echolocation signals that would be beneficial for swarm hunting.

Hunting on swarms rather than single insects could explain the presence of tiny insects in the diet of bats that can only be detected from very short distances ([8]). Swarm hunting might also shed light on early echolocating bats. According to our current fossil records, at about 52 million years ago, the first known bats (*Onychonycteris finneyi*) were able to fly [9]. Although we do not know if these early bats could echolocate, more and more evidence suggests that echolocation was an early trait. Recently, Wang et al. [10] used patterns of cochlear development to support the view that laryngeal echolocation is ancestral in bats. Arguably, access to the food source offered by nocturnal, flying insects was the adaptive advantage associated with flight and echolocation [11]. However, it is unclear how bats with a newly evolved and probably still rudimentary form of bio-sonar, could detect small insects, a task difficult even for modern human-engineered systems.

This riddle intensifies when taking into account the high costs of flight. Modern bats offset the costs of flight by eating 50% - 100% of their body weight every day [12]. Furthermore, these bats digest food in flight to immediately power flight [13]. Echolocation itself may be energetically inexpensive because of synchronization of search phase echolocation calls with wing downstrokes [14].

Most insectivorous bats are small animals (<50 g adult mass) which limits the size of their prey, thus requiring high rates of capture to maintain an energetic balance. However, because of the strong attenuation of high frequency sound waves in air, even modern bats, with echolocation strategies that evolved over millions of years can only detect a small insect from a very short range (<10m[7]). It was thus probably extremely difficult for early bats to detect small insects with their rudimentary echolocation, and it is therefore hard to imagine how these ancient bats could supply the high energetic demands dictated by flight.

We propose that detecting and hunting on swarms of insects rather than on individual prey could have facilitated a gradual improvement in the insect-detection abilities of bats. During evolution bats probably improved their ability to detect smaller swarms and eventually also single small insects. Even today, many bats hunt on concentrations of insects ([15]) such as mating swarms ([16], [17], [18]) or insects gathering at lights ([19], [20], [21]) and some species appear to prefer insect swarms [21]. Some bats even seem to have evolved special swarm-hunting strategies: e.g., *Pipistrellus abramus* appears to track and close-in on one flying insect, while tracking the next one (Fujioka et al. 2016[22]; Sumiya et al 2017[23]).

We hypothesized that many insectivorous bats are specialized for foraging on concentrations of insects. This generates two specific predictions: (1) Detecting concentrations of prey increases a bat’s effective hunting range. (2) The echolocation signal of some bats has evolved to suit swarm detection.

## Results

We generated 3D computer models of swarms and used both analytical calculations and acoustic Finite-Boundary-Element simulations (BEMFA; Boundary Element Method For Acoustics, [24]) to assess insect swarm echoes. In these models, we used two types of reflectors. Point reflectors allowed us to assess the effect of relative target positions in a swarm. Three dimensional mosquito-like reflectors allowed us to assess the effect of both individual insects in the swarm and the global swarm structure. We verified our computer simulations by recording echoes of real physical beads arranged like an insect swarm. Then we recorded echoes from an actual midge swarm in the wild. We tested different swarm structures, but all could be characterized by two parameters: the number of insects in the swarm (N) and the average distance between neighbors in the swarm (R). Each measurement we present is based on 100 realizations of random swarms with specific parameters (i.e., a specific combination of R and N). The exact positions of the insects in each realization were randomized by distributing them around centers R mm apart based on a Gaussian distribution (Methods). This ensured that no two swarms we examined were identical (for an example, see Figure 1A). For all of the analyses, we used a sonar signal covering a frequency range of 12-80 kHz, with equal intensity in all frequencies (Methods).

**Figure 1.**
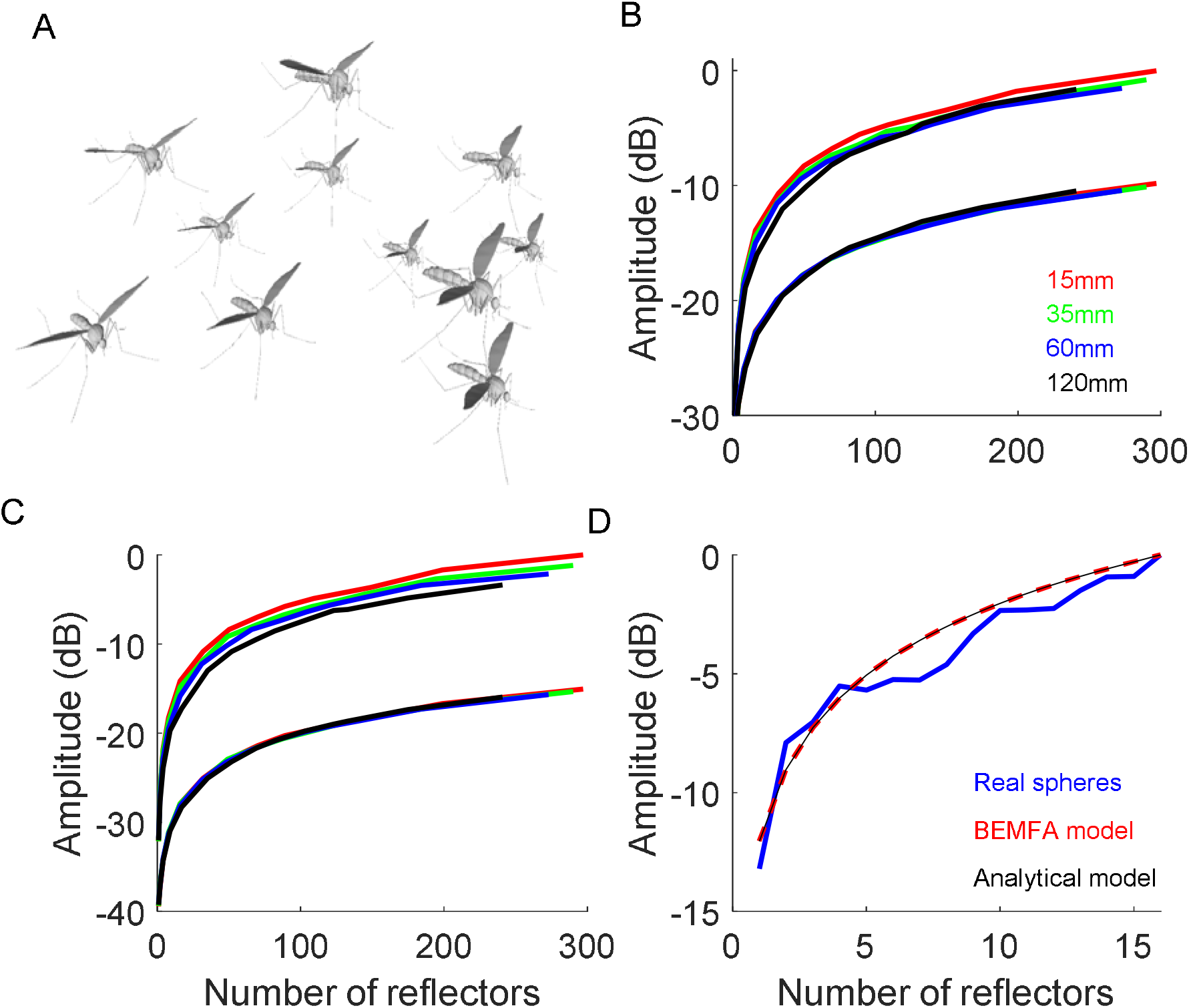
Swarm echo intensity increases with number of insects. (A) An illustration of one 3D realization of a mosquito swarm. (B-D) Echo intensity as function of the number of insects for different typical inter-insect distances (R’s) presented in different colors. Panels B-C show two curves for each color – one with the average and one with the maximum intensity (mean and maximum absolute pressure in dB). We normalized the maximum pressure to zero dB (and measure everything else in dB relative to max). Each point in the graphs is an average of 100 swarm realizations. (B) Results for the analytic model with point reflector. (C) Results for the analytic simulation with mosquito reflectors. In both B-C the higher curves represent the peak-to-peak intensities while the lower ones represent the average intensities. (D) Results for the physical beads (blue) and for the BEMFA simulation (red dashed). The thin black line shows the theoretical 3dB increase with doubling of reflector number.

### Intensity rise with swarm-size

We found that echoes of bat echolocation calls returning from swarms of insects enhanced the probability of prey-echo detection by bats. Every doubling of the number of insects covered by a single bio-sonar beam increased echo intensity by 3dB. This was true for point reflectors (Figure 1B) and mosquito-shaped reflectors (Figure 1C). The result was independent of the mosquito-orientation and bat view-angle (Figure S-1 and S-2). We validated this result for real bead echoes (Figure 1D). The physics of waves predict this increase which results from a summation of non-coherent signals [25]. The average distance between insects in a swarm had a negligible effect on echo intensity (Figure 1). Note that in our simulation, the echolocation beam is omnidirectional, invariably covering the swarm, and thus the entire swarm was always covered by the beam (Methods). This is mostly true in reality too, but occasionally if the bat comes very close to the swarm, or if the swarm is very large, the echolocation beam might only cover part of the swarm and only this part will contribute to echo intensity. In the mosquito-swarm, the absolute intensity of the echoes depended on the size of the specific insect. Swarms of larger insects would shift the entire graph upwards, but the 3dB increase per doubling of the number of insects would hold for insects of any size. Mixed swarms (of different insect species) would behave the same, according to the average insect size.

On average, an echo from a swarm increased by 3dB as a function of the number of reflectors independently of the echolocation signal and the structure of the swarm. However, occasionally, the echo strength of a given swarm could also be *reduced* by the interference generated by multiple echoes. We found that for each realization of the swarm, peak echo intensity occurred at completely different frequencies depending on the distances between the receiver and each reflector (Figure 2, note how the two - blue and red - patterns in each panel are completely different). The exact frequencies where spectral peaks and notches occur in a given swarm echo, depend on the specific interference pattern generated by every realization of the swarm. In a real swarm, the frequency producing the most intense echo is impossible to predict a priori as the insects in the swarm are constantly changing positions. Therefore, in the worst case, a bat using a pure tone (or a narrowband echolocation signal) might receive an echo that is more than 15dB weaker than the most intense echo it could have received from the same swarm, had it used signals of a different frequency (compare the dramatic intensity difference between two nearby frequencies in an exemplar swarm realization depicted by two black arrows Fig. 2B, left and right). Figure 2 shows that the spectrum changes more slowly when the inter-target distance is small (R small; compare upper and lower panels of each pair). This effect is quantified for all swarm densities and number of targets in the supplementary materials (Figure S3-4), showing that trough-width increases with decreasing inter-target distance.

**Figure 2.**
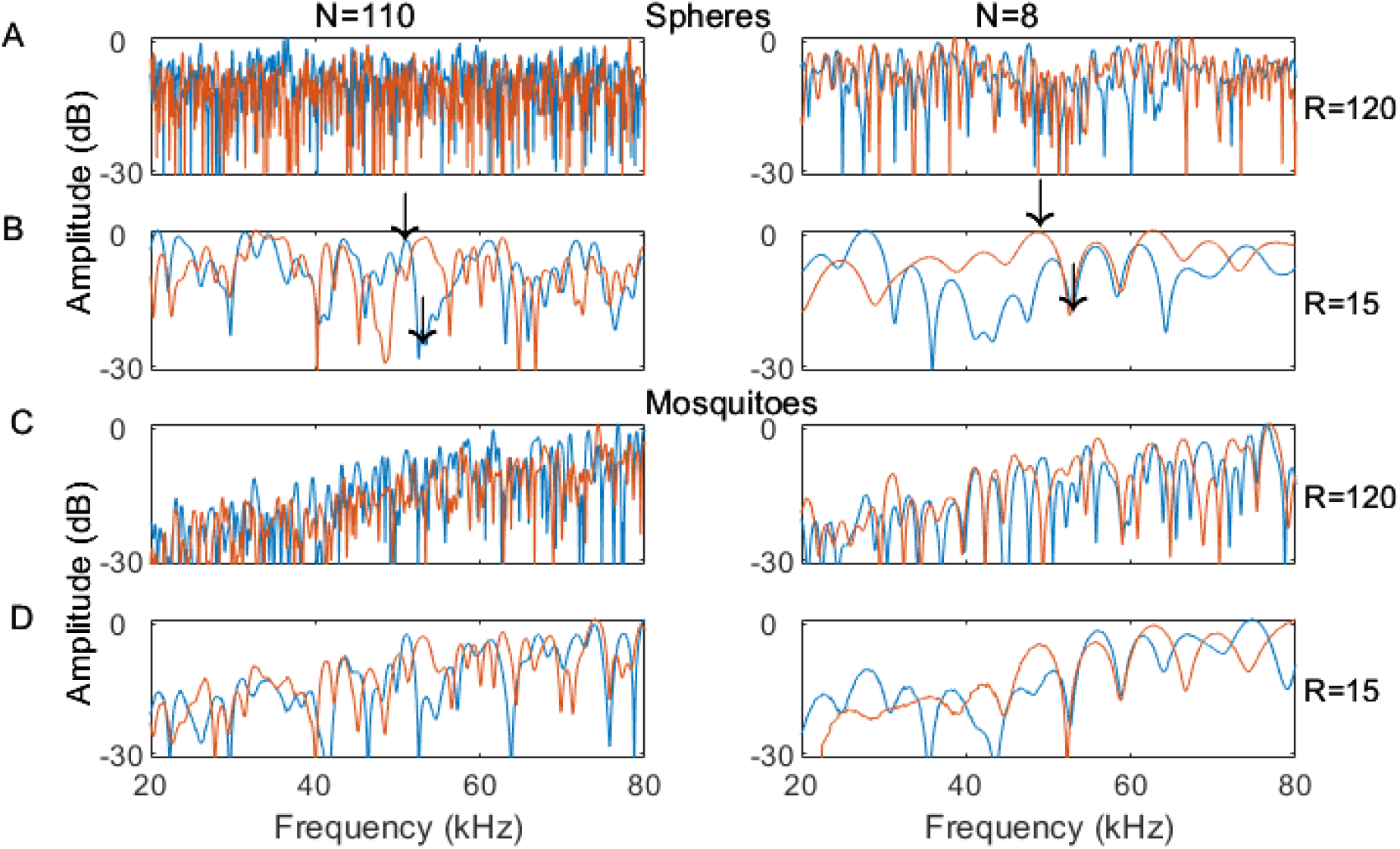
The spectrum of a swarm echo is highly stochastic. The spectra of four different swarm structures are presented for both point reflectors (four top panels) and mosquito reflectors (four bottom panels). All examples in the left column have N=110 reflectors while examples in the right column have N=8 reflectors. Panels A and C (rows 1 and 3) represent swarms with an inter-reflector distance of R=120mm while panels B and E (rows 2 and 4) represent swarms with an inter-reflector distance of R=15mm. Two realizations (orange and blue lines) are presented for each combination of parameters. Note how echo intensity (Y-axis) is strongly but stochastically frequency dependent (X-axis), and how two echoes returning from the same structure (compare pairs where N and R are identical) can have completely different spectra. We validated that the spectrum of an analytically simulated echo and a real echo returning from a group of beads with the same spatial structure are very similar (Figures S6-8). The mean intensity of the mosquito spectra increases continuously with frequency. This is due to the relation between the sound wavelength and the size of the mosquito. For the point reflectors (left) we did not model this effect because we aimed to focus on the role of swarm structure only. Note that the emitted spectrum is flat (each frequency has the same energy) so the differences observed in the reflected spectrum result mostly interference (and also from the frequency dependent atmospheric attenuation).

### Estimating the optimal bandwidth for detection of insect swarms

Because of the strong but stochastic dependency of echo intensity on the frequency of the echolocation signal, we estimated what bandwidth would be sufficient to guarantee high echo intensity for *any* given swarm structure. The average echo intensity improved with bandwidth, but this improvement leveled-out beyond a 6-10 kHz bandwidth (Figure 3). A bandwidth of 6-10 kHz should ensure that the bat always receives the maximal attainable echo intensity. This bandwidth is independent of the frequency of the echolocation signal (Figure 3). On average, a bat using a pure tone signal would receive swarm-echoes that are ~8dB weaker than when using a signal with 6-10 kHz bandwidth (Figure 3) and in extreme situations the echoes can be as much as 15-20dB weaker (Supplementary Figure S-5). Note that, as with the 3dB intensity gain function, the orientation of the simulated insects in a swarm and bat view-angle do not alter the relationship between bandwidth and echo-gain as displayed in Figure 3 (see control simulations in Supplementary Figures S-1 and S-2.

**Figure 3.**
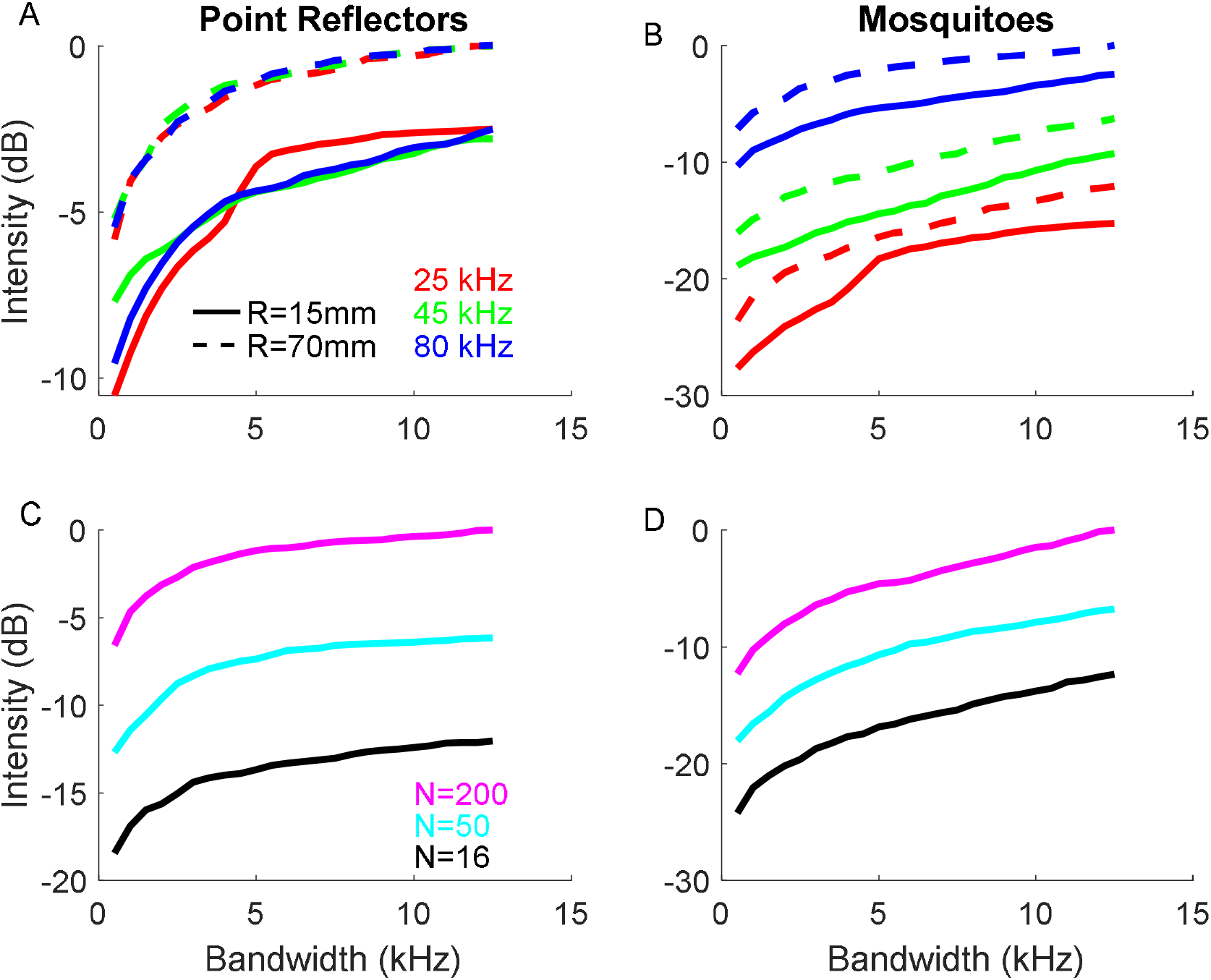
An echolocation signal with a bandwidth of a few kHz ensures high echo intensity. **(A)** Echo intensity as a function of bandwidth for a 100-reflector swarm with three different upper - frequencies (colors) and two different inter-reflector distances (R=15 and 120mm represented by solid vs. dashed lines respectively). Echoes of dense swarms (R=15) are slightly weaker than echoes of more spread swarms (R=120mm). This effect is caused by the spread swarm having a larger radius and extending closer to the bat, resulting in a louder echo. **(B)** The same as in A but for 100-mosquito-like reflectors; In both A-B the reported frequencies refer to highest frequency of the signal we used. **(C)** The echo intensity as a function of bandwidth for a point-reflector swarm with different numbers of reflectors (depicted by different colors). The highest frequency of the signals was 25 kHz for these simulations. **(D)** The same as in **(C)** but for mosquito-like reflectors. Each point (in all panels A-D) is based on generating 100 stochastic swarm realizations, calculating the loudest peak over the relevant bandwidth (depicted on the x-axis). Note that the lowest bandwidth is not 0 Hz, but 500 Hz.

So far, we have only addressed echo intensity. A bat’s actual success depends on the detection range of an object, which is a function of the intensity of the echoes but also the frequencies therein. Differential atmospheric attenuation means that echoes with the same intensity may be detectable at quite different distances. In general, lower frequencies attenuate less, and their echoes can be detected from longer distances. So we estimated the detection range as a function of the number of insects in the swarm for different frequencies. This analysis demonstrated the great advantage of using echolocation calls of lower frequencies to detect swarms (Figure 4). The advantage of low frequencies is clear for both mosquitoes and point reflectors. It is, however, less pronounced for mosquitoes because they reflect less at lower frequencies (when the wavelength is large relative to their size).

**Figure 4.**
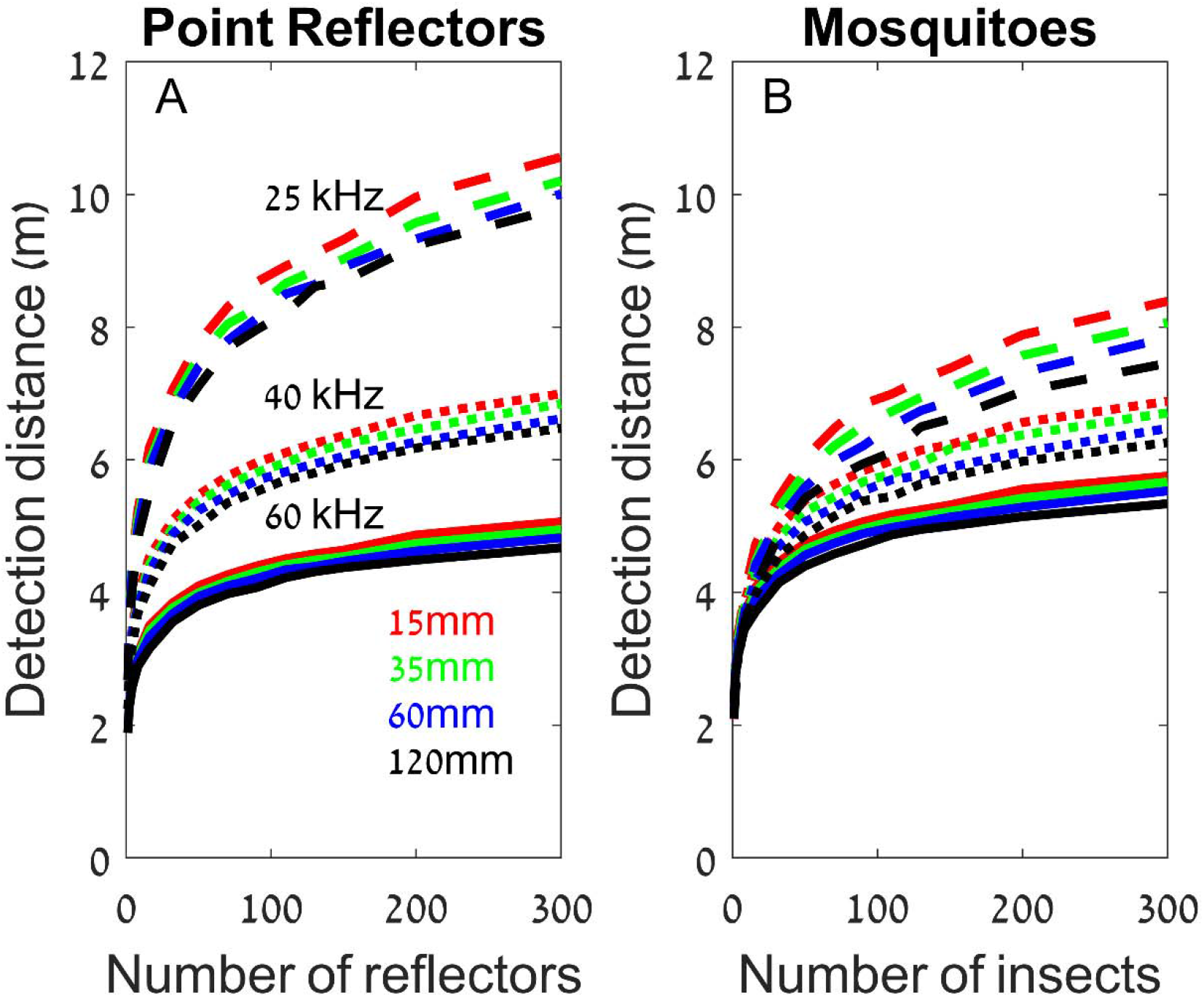
Swarms can be detected from larger distances than single targets. Detection range as a function of swarm size for a swarm of spheres (A) or mosquitoes (B). To create this figure we assumed an emission intensity of 130dB SPL (at 10cm from the mouth), a detection threshold of 10dB SPL and a target strength of −80dB at 1m (−60dB @10cm) for the spheres. The colors depict different inter-insect distances in the swarm and three types of lines represent different terminal frequencies: 25, 40 and 60 kHz (represented by dashed, dotted and solid lines respectively). Atmospheric attenuation used: 0.5dB/m; 1.2dB/m and 2.3dB/m, respectively. Data are the same as those used to plot figure 1.

In theory, bats could emit several pulses instead of broadening their bandwidth in order to improve their chances of detecting a swarm of insects (because one of these echoes would generate peak intensity). We examined this as well by testing how many signals would be required in order to match the detection of a single call with a certain bandwidth (Methods). Results demonstrate that a bat would need to emit more than 10 signals to match the detection probability of a single signal with 1.5kHz bandwidth (Figure S-9).

### Echo measurements of real insects

Finally, to confirm our prediction of enhanced detection through swarm hunting, we compared the detectability of a single midge and an actual swarm of midges (*Culex pipiens*). To this end, we used a bio-mimetic sonar system (with an pulse of: 1ms 55-23 kHz and 100-50 kHz) to record echoes of the swarm in the field and to record echoes of an individual midge in the lab from equal distance (the single insect was tethered on a 0.1mm wire in our acoustic room, Methods). The peak intensity of the swarm echo was on average 12.8 (+/− 2.3) dB (n=77) louder than that of the single insect echo (Figure 5). The swarm echo was also ca. twice longer (because echoes continued arriving from farther midges) thus allowing to accumulate echoes over a longer period, which would improve detection by the auditory system. Because the swarm was ensonified in the field, we cannot estimate the accurate number of midges that reflected echoes, but as our artificial beam was narrower than that of actual bats, a real bat would probably gain even more from searching for a swarm because its wide beam would result in reflections from more insects.

**Figure 5.**
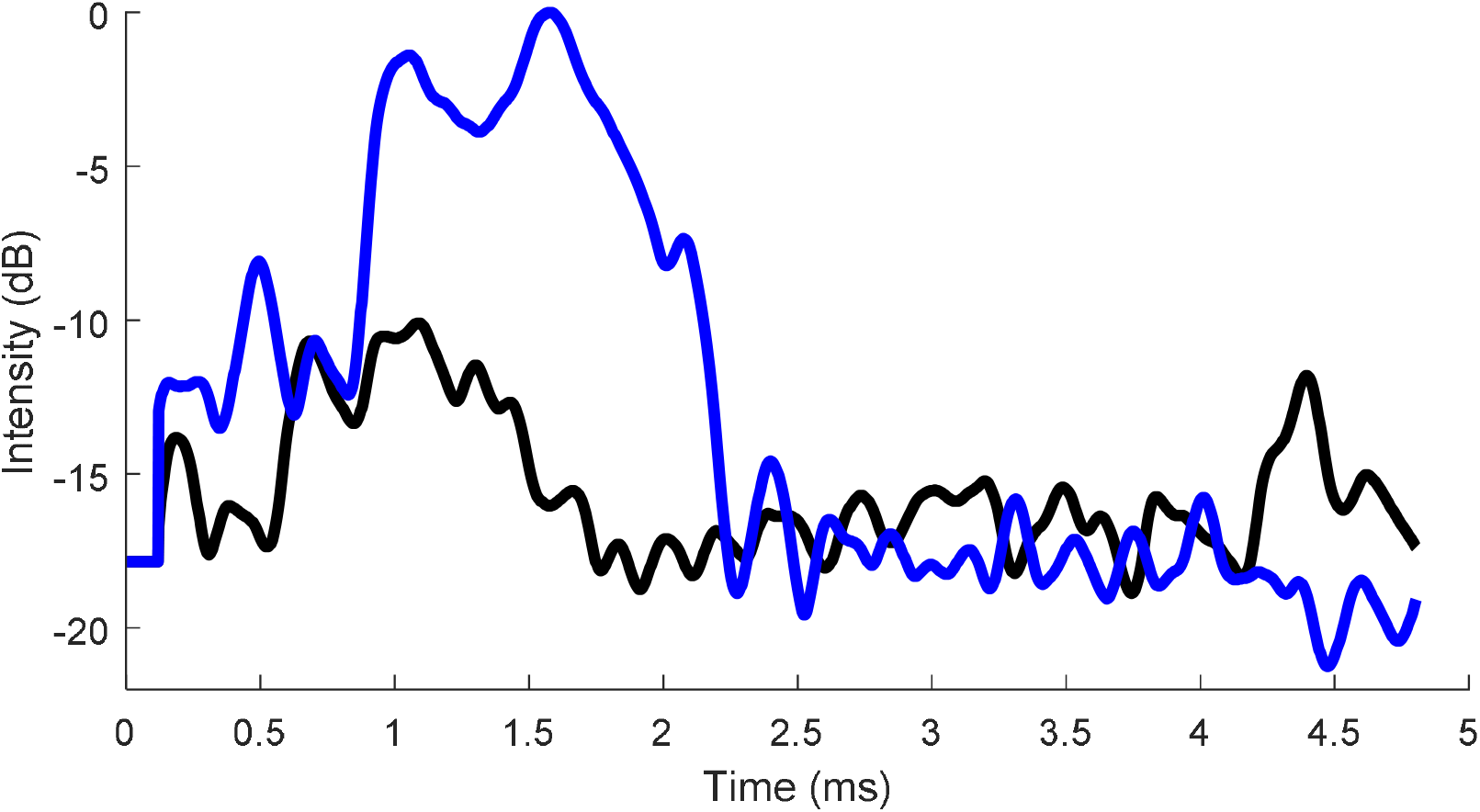
A comparison of the echoes of one midge (black) and a midge swarm (blue). The envelope of a one swarm and one single insect echoes are presented.

## Discussion

Our data support the prediction that swarm hunting increases prey detection range. Bats can use echolocation to detect swarms of insects at greater ranges than single insects. Echoes from a swarm of insects could serve as ‘beacons’ attracting bats from larger distances. A bat using echolocation calls with most energy at 25 kHz emitted at 130 dB SPL re 10 cm could detect a swarm of 300 midges from ~8 m compared to only ~2 m for a single midge. We support our theoretical calculations with recordings of echoes from actual swarms in the field. Interestingly, the interference of the sound waves, reflected from a swarm of insects means that echoes from swarms can sometimes be much weaker than those from single insects. Echolocating bats can overcome this problem by producing signals of broader bandwidth. A bandwidth of 6-10 kHz would guarantee the loudest possible echo from any angle relative to the swarm. Moreover, using echolocation calls dominated by lower frequencies (around 25 kHz) would be most suitable for detecting swarms. This analysis only considered the gain from echo intensity. Echoes from a swarm of insects will also be longer than those of a single insect because reflectors are spread over a large volume (compare blue to black in Figure 5). The integration time of the hearing system would further increase detectability of such longer echoes by up to a few more dBs (see Methods).

Interestingly, many bats that specialize in searching for prey in open space (i.e., far from nearby objects) use very shallow FM echolocation signals, which are characterized by peak frequencies ranging between 20-40 kHz with a bandwidth of 3-8 kHz (Table 1). This bandwidth is probably not a result of an inability to maintain a constant frequency. Even humans can control the pitch of their voice more accurately than 1% (down to 0.2%) over more than 5 secs (own measurements). To stay within 500 Hz, -our “zero bandwidth” point- (Figure 3), should therefore be easily accomplished by bats. Sweeps over several kHz, as displayed in Table 1 must therefore be produced deliberately.

**Table 1.**
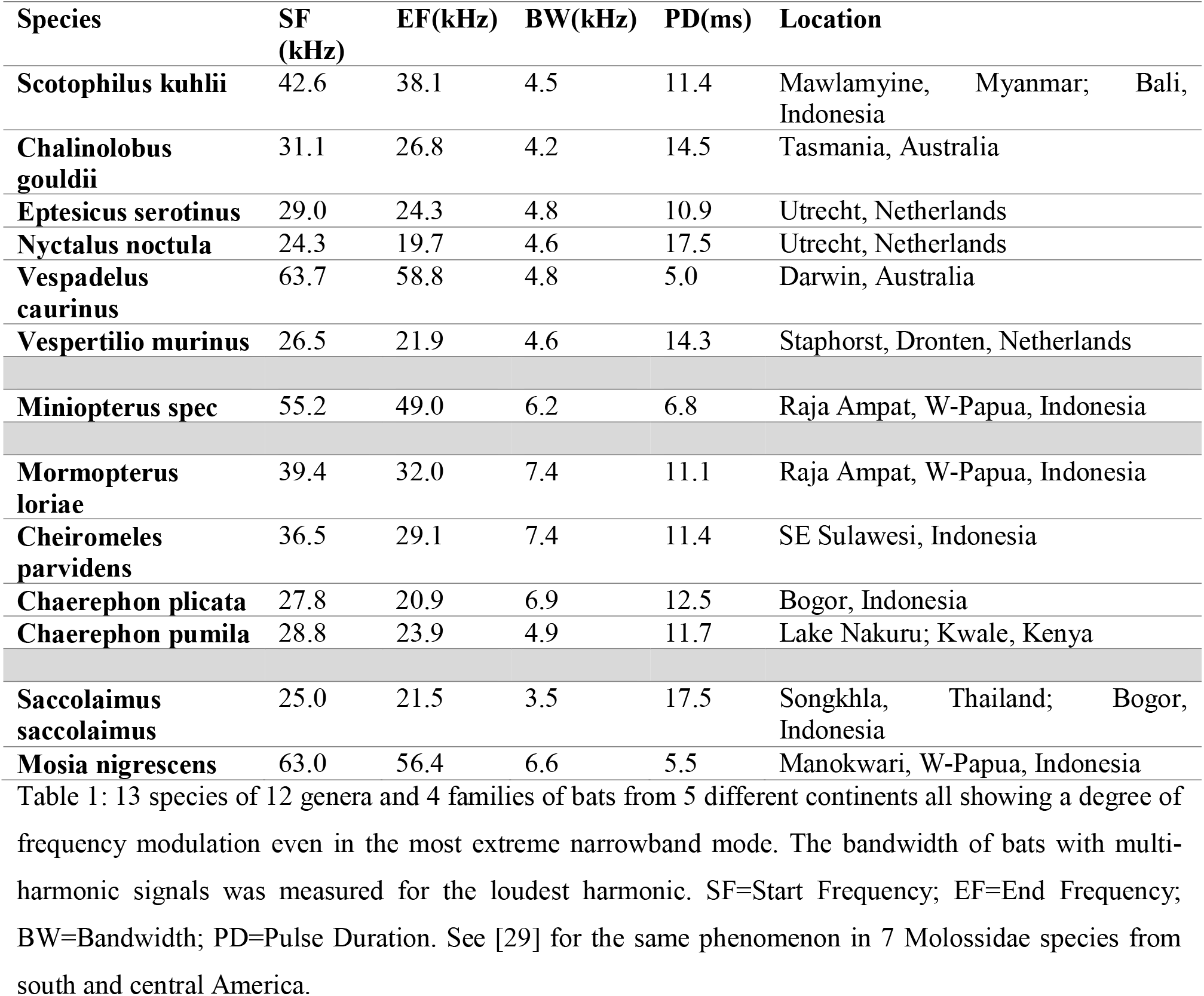
13 species of 12 genera and 4 families of bats from 5 different continents all showing a degree of frequency modulation even in the most extreme narrowband mode. The bandwidth of bats with multiharmonic signals was measured for the loudest harmonic. SF=Start Frequency; EF=End Frequency; BW=Bandwidth; PD=Pulse Duration. See [29] for the same phenomenon in 7 Molossidae species from south and central America.

Apart from modulating the constant frequency, another strategy, used by many bats species, is to alternate between signals with terminal frequencies several kHz apart (typically 3-10 kHz) [26], [27], observed particularly when bats hunt for prey [28]. This alternation would effectively be the same as using a signal with a bandwidth of several kHz.

We show that increasing signal bandwidth is beneficial when searching for insect swarms. However, increasing frequency bandwidth comes at a cost. In fact, when detecting a single insect with a mammalian hearing system, allocating more energy to less frequencies (i.e., less bandwidth) would be beneficial for detection. The exact bandwidth used by a bat species when searching for prey could thus be a trade-off between two needs: allocating energy to less frequencies to improve Signal-to-Noise-Ratio, and dividing energy between many frequencies to overcome echo-notches resulting from interference of multiple swarm echoes. The most suitable peak frequency might also be different for detecting a single insect vs. a swarm (for comparison [5], [6]). Note, that we are only considering detection here. When a prey has already been detected, and localization is at stake, bandwidth is important for localization as well.

The advent of DNA barcode analysis has made it possible to determine which species of insects are eaten by bats (e.g. [30], [31], [8]). Such bat diet analyses reveal that bats using low frequency signals eat mosquitos with wing lengths below 6.5 mm in large numbers that would be almost impossible to detect using bio-sonar because: (1) The target strength of these insects would be around −73 dB or lower at 25 kHz, [6], [7] and (2) The bat would have to detect them from over 2m because of the overlap free window resulting from the pulse durations ([32]). Our results show that such small insects can be detected when occurring in large swarms, which many families of insects exhibit (e.g., *Chironomidae, Blissidae, Noctuidae, Hydropsychidae, Formicidae*), due to active gathering or wind. A swarm of 200 small insects compact enough to be covered by the echolocation beam has an echo intensity equivalent to that of a large (>1.5cm wing-length) moth that can be easily detected by a bat (target strength −57 dB @1m). It is therefore likely that low frequency bats only detect very small insects when the insects aggregate in swarms.

In theory, bats could emit multiple signals instead of increasing signal bandwidth, but we show that this strategy would require many signals, and would thus result in a waste of time and perhaps also energy (if the wingbeat to call synchronization breaks).

Our results show that the randomness of the spectrum due to swarming-targets depends little on the shape of each of the elements (point targets vs mosquitoes in various orientations). The noise introduced by the constellation most likely overrides the randomness in the echoes of each of the individual targets.

Hunting for swarms could thus increase both the chances of finding food and the variety of consumed food items, enabling bats to detect and feed on small insects. Swarm hunting could thus explain how echolocation has evolved to enable insect hunting. It is believed that the earliest bats were insectivores, but the detection of small single insects using bio-sonar is extremely difficult today, and was probably much more difficult with rudimentary echolocation and rudimentary brain processing of echolocation. One possibility is that early bats only ate very large insects, but that would strongly limit their diet. We show that the tendency of insects to aggregate in swarms facilitates the detection of small insects, which would have substantially broadened the diet of early bats. Swarm hunting could thus have mediated the evolutionary transition to more sophisticated echolocation systems. Detecting smaller and smaller swarms of insects down to detecting single insects of small size could have made this transition infinitely smooth.

## Methods

### Generating point target swarms

As flocking, swarm formation- and movement have become a popular topic in biology (e.g. [33]), a number of detailed studies and modeling efforts to copy natural behavior of mosquito swarms already exist ([34], [35]). Following these studies, we generated point reflector swarms by dividing space into 3D cells (voxels) and placing a virtual point reflector at the center of the cell with Gaussian noise. We controlled two parameters of the swarm, the number of targets (N) and the distances between cell centers (R) which were based on inter-target distances observed in natural insect swarms (N’s between 10 to 100; R’s between 35 to 130mm) (e.g. [31, 33]). To cover different types of swarms encountered in nature, we simulated insect swarms with average nearest neighbor distances R of 15, 35, 60 and 120mm. For each task we simulated 100 swarms. Each swarm realization differed from the other 99 realizations due to the Gaussian noise in the placement of the targets in the centers of the cells.

The standard deviations of these position distributions were: 6; 15; 26 and 52mm (for each of the above R’s respectively).

In this paper, we will refer to a swarm realization as a newly generated swarm consisting of the same number of targets, but in totally new positions while still having the same statistical properties (i.e. same N and R). When taking into account the inter-insect distance (R) and the number of insects, our simulated swarms had overall diameters of 0.13; 0.31; 0.52 and 1m for our swarms of R=15; 35; 60 and 120mm respectively (for N=300). We generated swarms with the following sizes: 1, 2, 4, 8, 16, 32, 50, 70, 90, 110, 130, 150, 200 and 300 targets. The diameter of any of our swarms declined with decreasing the number of insects similar to what was reported for Chironomid-clouds (Kelley and Ouellette 2013 [34]).

### Analytic calculation of point-target swarm spectrum

The echo intensity (at any frequency f) from any realization of a random point cloud, consisting of N targets was calculated as:

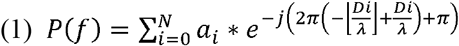

Where Di is the 2 way distance from emitter/receiver to point target i, a_i_ is the spherical loss factor for each target i, λ is the wavelength and −j denotes the imaginary part of a complex number (j^2=−1). Square brackets denote rounding to floor (obtaining the integer). The factor a_i_ denotes the 6dB loss in pressure with doubling of distance relative to 0.1m from the target (where target strength was set to −60dB) on a linear pressure scale. This equation does not specify atmospheric attenuation. In calculations where atmospheric attenuation becomes relevant (i.e., Figure 4) we added this effect using Bass et al. (1995)[37]. We used both boundary element model simulations and actual physical models to validate this analytical method, see supplementary results (Figures S6-8).

### Estimating the echoes of insect swarms using the analytic model

In the case of the point targets, the echoes of each point were not distorted in phase or amplitude. This enabled us to study the effect of swarm’s structure independently of the response of its constituents. However, natural swarms are composed of insects that reflect echoes that may shift in phase and have a specific frequency response. We therefore measured the complex spectrum of a very detailed 3D mesh model of a mosquito including legs, antenna, eyes, segmented body and wings (between 12-80 kHz, see next section) from every angle with steps of 5 degrees. Next, we used this dataset as a lookup table to generate the echo of an insect from any desired orientation, or in our case; to design a swarm of insects each in any desired orientation. The insects were positioned in space just like the point reflectors (see above) but we also had to determine their orientation. In real insect swarms, individuals tend to align the orientation of their bodies. To mimic this, all insects were horizontal relative to earth facing the same direction. Since in reality insects are not always perfectly aligned, we mimicked a swarm in which yaw- and roll-angle varied probabilistically with +/− 30 degrees. The bat was facing the center of the swarm (2m from the center) viewing it from above with an angle almost perpendicular to the wings (−15 deg relative to the dorsal/ventral axis) so that the wings produce loud echoes. To verify the robustness of the data generated with this type of swarm, we also carried out control-simulations, one in which the swarm was ‘seen’ laterally (same +/− degree Gaussian spread) and one in which the alignment of the insects within the swarm was completely randomized.

Finally, in order to estimate the insect-swarm echo, we used equation 1, the spectrum was multiplied by the insect’s complex frequency response for the specific angle (obtained from the mesh model simulations).

### Estimating the echo of an insect mesh using Boundary Element Model Simulations (BEMFA)

To estimate the angle dependent insect frequency response, we used a 3D mesh model of a mosquito. The model was complete with eyes, antennae, legs, wings and body segmentation.

Using Meshlab [38], we checked our mesh model for any holes, non-manifold elements, face coherence, unreferenced vertices and we verified that all normal vectors were directed out of the body into “air”. The longest element size was still 26 times smaller than the smallest wavelength used (at 80 kHz) in our simulation. Boundary element simulations were carried out with the Matlab routine Bemfa [24]. Bemfa uses Green’s function to solve linear differential equations of acoustic wave propagation from source to receiver by convolving a boundary condition with a transfer function. The result is a steady-state solution of the acoustic reflections in the frequency-domain of the mesh-object. Bemfa relies on the CHIEF-method [39] to deal with irregular frequencies with undetermined solution. This means that in order to solve the equation from mesh-reflectors to receiver, CHIEF points are placed randomly inside the mesh volume where the sound pressure is zero. From this, a second set of linear equations is derived that do lead to a unique solution. We checked meticulously that the CHIEF-points we generated in each simulation were always placed inside the insect-body (including wings and legs) as our routine was supposed to place them. We tested this routine by generating an excess number of CHIEF-points several times with the sole purpose of verifying their positions. In the actual simulations we used 4% CHIEF-points relative to face count as the object had a very contorted structure. The mosquito was scaled such that each wing was 2.7mm long from wingtip to wing-base and it was placed at 20 cm from a virtual point sound emitting source. Sound pressure level at the location of the insect mesh was 86 dB SPL. Emission level is the same for all frequencies (12 to 80 kHz). The point source is rotated around the insect in an entire hemisphere with a radius of 0.2m from the target in steps of 5 degrees in azimuth and elevation (echoes were measured at the same position of the source).

### Echo Analyses

#### Intensity rise with swarm-size

We calculated the echo spectrum for each target constellation between 12-80 kHz. From each spectrum we calculated the peak and average pressure (in dB), and took the average over 100 swarm realizations per condition (e.g., number of targets) and plotted the result against the number of point targets.

#### Estimating the optimal bandwidth for detection of insect swarms

For three frequencies (25, 45 and 80 kHz) we increased bandwidth with steps of 500 Hz down to 12.5 kHz below these upper frequencies, and calculated the maximum echo pressure for each echo (note that these maxima could occur at any frequency). For each bandwidth, we generated 100 simulations (each with a different stochastic swarm arrangement) from which we selected the 25 simulations that had the lowest maximum pressure values. We then plotted the mean of the maximum pressure value for these 25 simulations against bandwidth indicating that the bat could ensure receiving an echo with this intensity (or more) in 75% of the time. We also tested a more conservative value of 90% probability, following the same procedure and plotted the result in figure S-5.

### Ensonifications of bead swarms in the lab

#### Model cross-validation with a constellation of real objects

In order to evaluate the different simulation tools used in this paper, we also created a point cloud in real life, using wooden spherical beads. We also simulated this exact same cloud (of different sizes N) mathematically, using the same methods that were used to generate the simulations of insect swarms in this paper. We then compared the results of our simulations with each other and with recordings of real echoes of the real beads. To measure the echoes of real sphere constellations (n=1…16), we suspended wooden beads (2cm diameter) on 0.1mm diameter cloth strings in an acoustic chamber. Prior to ensonification, 2 IR reflectors (semi-sphere, 6mm diameter, 3X3 Designs Corp.) were placed on opposite sides of each bead. A system of 20 tracking cameras (a Motion - Analysis Corp system. 16 Raptor E cameras 1280 x 1024 pixels cameras and 4 Raptor-12 4096 x 3072 pixels cameras) was used to measure the exact locations (x,y,z) of each bead. The accuracy of this camera system was proven to have accuracy of ~1mm in an extensive series of performance- and verification tests [40]. The loudspeaker and microphone (used in the ensonification, see next section) were also localized by the camera system so that distance D from loudspeaker to each sphere could be calculated. The cloud of 16 randomly positioned spheres ranged from 0.93m to 1.18m distance from the speaker center (0.09m spread in height; 0.15m spread in width). Their average nearest neighbor distance (R) was 81.9mm within the range of swarms that we simulated.

A series of 15 narrowband sound pulses (3.5ms each) together covering the spectrum of 12 to 80 kHz were emitted at the spheres using the Avisoft Vifa loudspeaker (connected to an Avisoft 116 Player D/A). The signals were 15 linear frequency modulated sweeps with a bandwidth increasing from 4 to 14 kHz around a center frequency increasing exponentially from 12.2 to 76.2 kHz from sweep 1 to sweep 15. This signal design kept the wavelength change of the emitted pulse below a factor 1.8 for nearly all signals. Keeping the wavelength-change constant ensures that the potential echo-information is spread equally across channels. The echoes were recorded using an Avisoft CM16 condensator microphone (connected to an Hm116 A/D, Avisoft), attached as close as possible to the loudspeaker. This was repeated 9 times. Then the microphone was placed in the middle of the sphere-cloud, the spheres were removed and the same sound was recorded 9 times in order to estimate the emission’s frequency response and compensate for it (see below). Both paradigms were done with neutral phase and opposite phase (the signal but with a 180 phase difference). For every condition mentioned above the 9 spectra for each frequency band were averaged. The 15 averaged spectra were then merged from 12 to 80 kHz. This was done for the microphone in both positions. The final spectra from both positions were normalized and the ratio calculated between both spectra. This ratio is the difference between the incoming sound to the spheres and the sound reflected from the spheres. It is therefore the same as the echo of the spheres if a flat spectrum were used (as was the case in the analytic equations and the BEMFA simulations).

### Echo measurements of real insects in field

Echoes of wild mosquito swarms (*Culex pipiens*) were recorded at the brackish water inlets of Goeree Overflakkee, Netherlands, GPS 51.8038N 3.9649E. We used the same equipment as described under “Echo measurements”. The emitted pulses were 1ms sweeping from 55 to 23 kHz and from 100 to 50 kHz. We compared the echoes of the swarm with echoes from a single tethered insect (0.1mm wire diameter) suspended in an anechoic room at 50cm distance from the loudspeaker and microphone. We verified from the time between emission and echo in the recording that in the real insect swarm, mosquitoes were present in front of the loudspeaker at the distance used to ensonify the single insect in the lab, thus enabling us to compare echo strengths (Figure 5). We estimate the number of insects inside the beam between 4-20 insects. Note that many investigated bats appeared to have sound beams (−6dB beamwidth) of around 75 degrees (30-55 kHz) ([41]), whereas the Avisoft Vifa loudspeaker we used has a 19 and 12 degrees −6dB beamwidth for the following frequencies 30 and 55 kHz respectively (own measurements). Due to their 5x wider beam, bats would cover more insects in the swarm, and would thus experience a more pronounced difference than the one depicted in figure 5.

### The effect of temporal integration

A swarm of insects contains targets much nearer and also much further from the bat relative to the center of the swarm. A bat echolocating a swarm will thus experience a longer echo than if it were only targeting a single insect. The exact echo integration mechanisms in the hearing of bats is unknown, especially for detection situations using very long pulse durations, as bats do not use such pulses under laboratory conditions. We therefore calculated the increase in gain a bat would experience if its integration mechanisms were unlimited and perfect. These calculations provide information on what bats could maximally gain from the elongation of its echo due to swarm depth.

In our calculations we assumed the bat to be emitting pulses of 25 kHz with the relatively short pulse duration of 8 ms. The assumption of a short pulse duration will boost the possible integration-gain even further. We calculated for each swarm density by what fraction our 8ms pulse was lengthened due to swarm diameter. Following the LIEFTS model established by [42] we assumed a 6dB gain with doubling of echo-duration. For the 15mm nearest neighbor distance of insects we found 8.9 ms as longest echo (N=300). Following [42] this translates as a gain of 6.02*log2(8.9/8)= 0.9dB or an increase of 17 cm in detection range at 4m distance. For the least dense swarm type investigated the gain was higher; from 8ms to 15ms; 5.5 dB leading to a lengthening of detection range of 1.15m for prey at 4m. However, it should be kept in mind that this gain is the best achievable by a bat with optimal swarm type and with the bat using relatively short pulse durations. In any other situation the gain due to temporal integration will thus be very small relative to all other factors put forward in this paper.

### The benefits of bandwidth vs pulse repetition

To investigate whether bats could simply emit more than one pulse to enhance swarm detection (instead of increasing bandwidth), we calculated the number of pulses that would be required to emit in order to match the detection performance of each bandwidth. We did so by examining the echo-intensity distribution for each signal bandwidth (we simulated 100 swarms per bandwidth). We then measured how often the echo intensity (for a given bandwidth) was above the −4dB point relative to the peak intensity for full bandwidth signal (i.e., a signal with 12.5 kHz bandwidth). This fraction represents the probability of an echo to be within 4dB from the maximum potential intensity. The inverse of this fraction represents the number of pulses required (for the given bandwidth) for the echo to within 4dB from the maximum potential intensity.

## Acknowledgements

We thank Nir Gov (Weizman Institute) for providing advice to simulate life-like insect swarms. We also thank Vladimir Tourbabin for providing the Bemfa code and guiding us how to use it properly. Ofri Eitan helped us in obtaining the exact sphere coordinates for the cross-validation experiment. We thank Stuart Newson for providing recordings of *Chalinolobus gouldii* from Tasmania and Jens Rydell for providing recordings of *Chaerephon pumila* from Kenya. This research was funded by ONRG project N62909-16-1-2133.

