## Supplementary figure S1 to 9 for "The benefits of insect-swarm hunting to echolocating bats, and its influence on the evolution of bat echolocation signals"

**Supplementary Material**

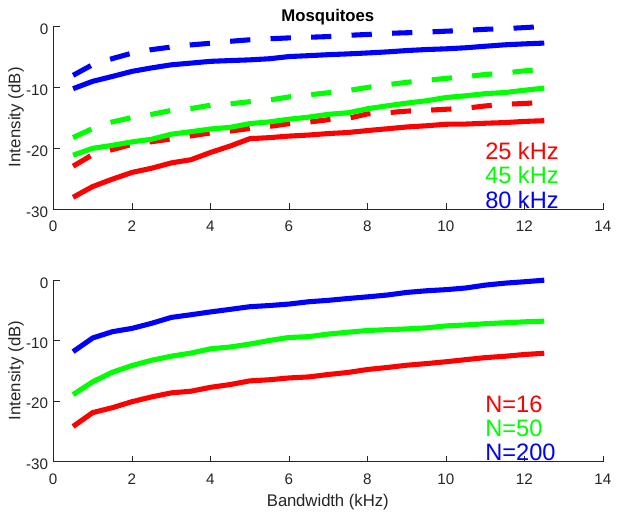

**Figure S1. An echolocation signal with a bandwidth of a few kHz ensures high echo intensity, same as Figure 3, but here with all insects oriented sideways, also with a +/-30 degrees Gaussian spread.** Upper graph: Solid line R=15mm, dashed line R=200mm, colors denote different frequencies. Lower graph: R=35mm, but with different swarm sizes. Results from this simulation reveal similar principles as in Figure 3 in the paper.

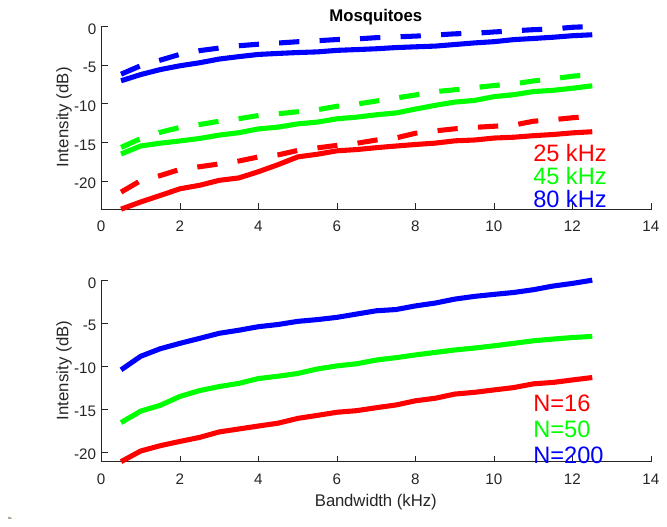

**Figure S2. An echolocation signal with a bandwidth of a few kHz ensures high echo intensity, same as Figure 3, but here with each insect oriented randomly.** Upper graph: Solid line R=15mm, dashed line R=200mm, colors denote different frequencies. Lower graph: R=35mm, but with different swarm sizes. Results from this simulation reveal similar principles as in Figure 3 in the paper.

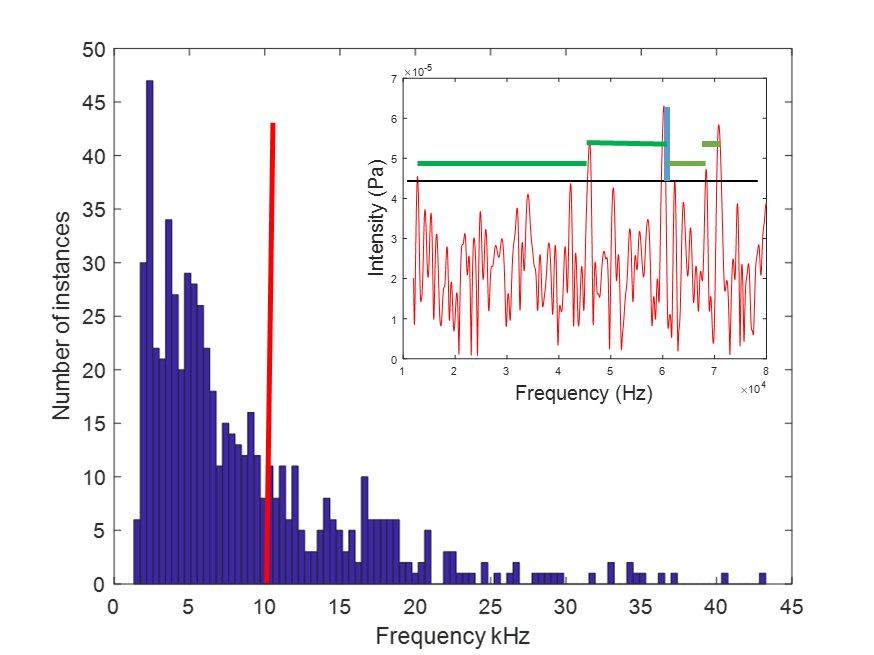

**Figure S3.** The distribution of spectral trough bandwidths (see top-right inset) for a swarm of 16 insects, and 100 trials swarm realizations (R=35mm). For each spectrum, the bandwidth of all troughs between pairs of peaks was measured. Peaks were defined as peaks that are no more than 20% lower than the maximum of the entire spectrum (black horizontal line inset; green horizontal lines: trough-widths). The bandwidth of a trough can be thought of as the minimal bandwidth allowing to receive a peak intensity for a given swarm realization. Red line shows that a bandwidth of 10 kHz would ensure that the bat receives the maximal echo (i.e., no more than 20% weaker than the absolute possible maximum) for 70% of the echoes (the line parts the histogram to 30:70%). We repeated this analysis for different peak criteria (10, 30, 40%) and the pattern is the same (the bandwidth would obviously change accordingly). We call this point of 70% BWc – the critical bandwidth.

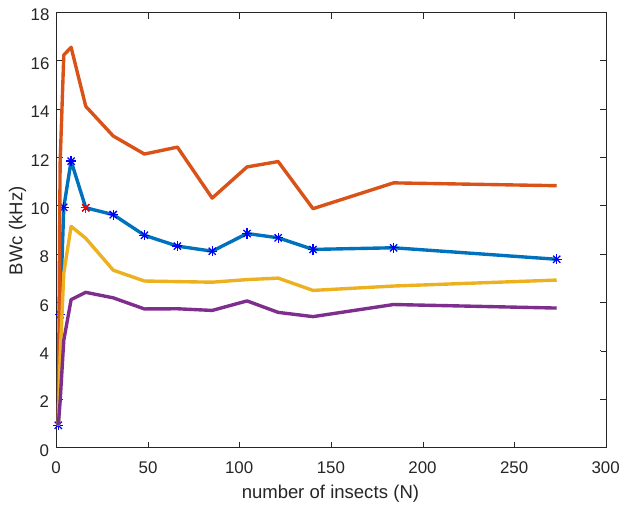

**Figure S4.** The critical bandwidth (BWc, see Figure S1) does not depend on the number of targets in the swarm (for more than 2 targets). BWc was extracted as in figure S1. Red star (on blue line) is the same point as the red line shown in figure S1. The same pattern was observed for swarms of different densities (R=15mm; R=60mm and 120mm; red, yellow and purple lines respectively). More bandwidth is needed for denser swarms (compare red and purple lines) and less bandwidth is needed to capture spectral peaks of sparser swarms. This can also be learned from Figure 2 in the main text.

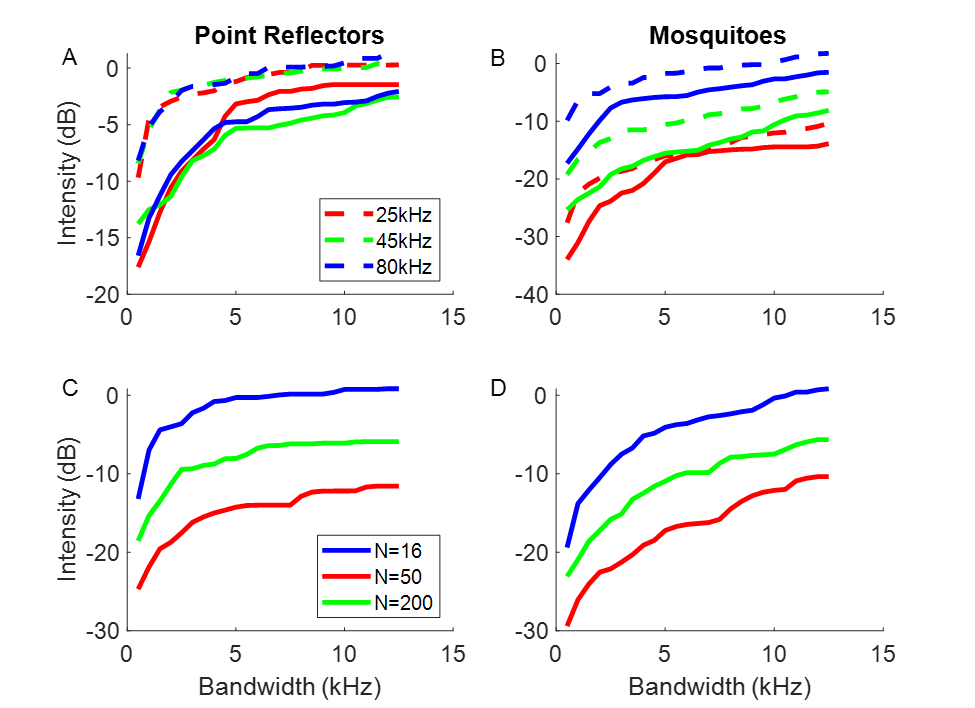

**Figure S5.** All details identical to figure 3 in the main text, except here we took the worst 10% cases of 100 simulations. Worst cases: in the minimal bandwidth case (500 Hz), a bat is faced with the worst spectral trough of 100 simulations. The figures show that adding bandwidth improves swarm detection by 15-20dB on average in this worst case scenario. (A) Echo intensity as a function of bandwidth for a 100-reflector swarm with three different upper -frequencies (colors) and two different inter-reflector distances (15 and 120mm represented by solid vs. dashed lines respectively). (B) The same as in (A) but for 100-mosquito-like reflectors. (C) The echo intensity as a function of bandwidth for a point-reflector swarm with different numbers of reflectors (depicted by different colors). The upper frequency was 25 kHz for these simulations. (D) The same as in (C) but for mosquito-like reflectors. Each point (in all panels A-D) is based on generating 100 stochastic swarm realizations, calculating the loudest peak over the relevant bandwidth (depicted on the x-axis). Note that the lowest bandwidth is not 0 Hz, but 500 Hz, therefore displaying only intended bandwidth and not natural variation (see Discussion).

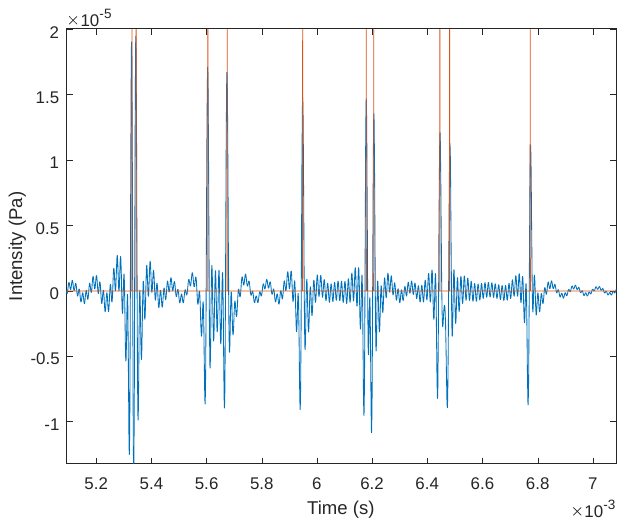

**Figure S6. Cross-validation of analytical-, simulation- and empirical methods.** We tested the validity of the analytic model (equation 1) numerically by generating swarms with 1 to 10 points at known distances D_i_, calculating the power spectrum of their echoes (using eq. 1), and then estimating the impulse response by means of an inverse Fourier transform. The impulse response preserves the temporal information which can be compared to the distances of the reflectors. The spectra were calculated between 12 to 80 kHz in steps of 100Hz. The figure shows the actual point distances D_i_ for a single swarm with 10-points (red) and the impulse response generated according to equation 1 (blue). The distances are presented over a time axis to ease the comparison.

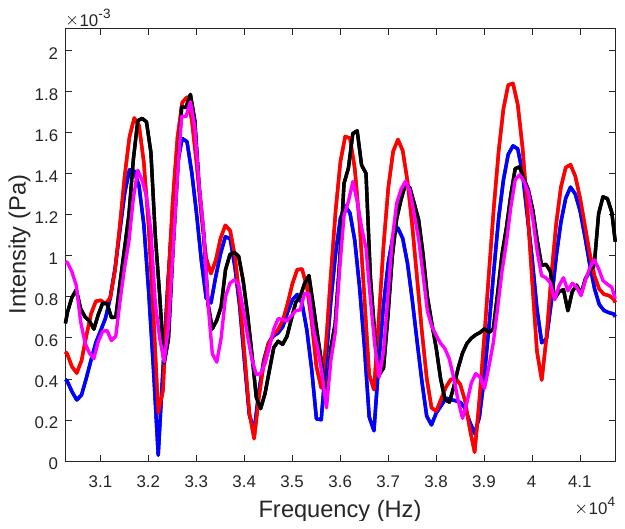

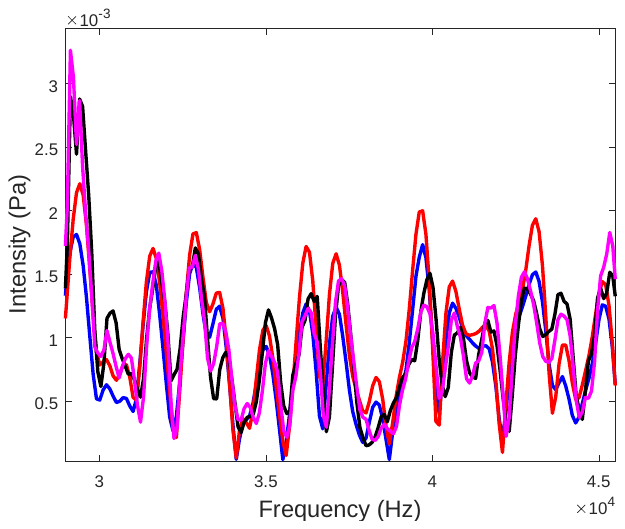

**Figure S7. Comparison of echo generation models.** The spectrum of three models is presented: the analytic model (red), the boundary element model (blue) and the actual recording of real beads (in black and in magenta for echoes for which the emission was shifted by 180 degrees). **Top**: the spectrum of a 7-reflector model is presented. Bottom: the spectrum of an 8-reflector model is presented. For a better comparison, the spectrum between 30-42 kHz is enlarged. Note that the real beads had a 2cm diameter which could explain the slight differences between the actual echoes and the simulated ones.

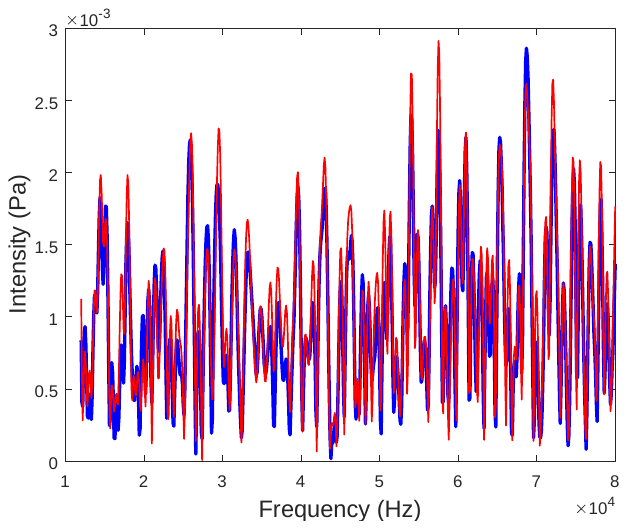

**Figure S8. Comparison of echo generation models.** The spectrum of two models is presented for a 10-reflector model: the analytic model (red), the boundary element model (blue). We do not show the data for the physical bead model because it would be difficult to see with another graph overlaid.

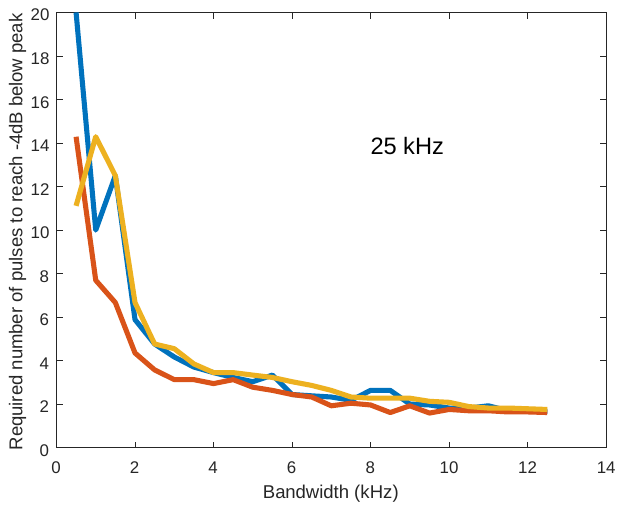

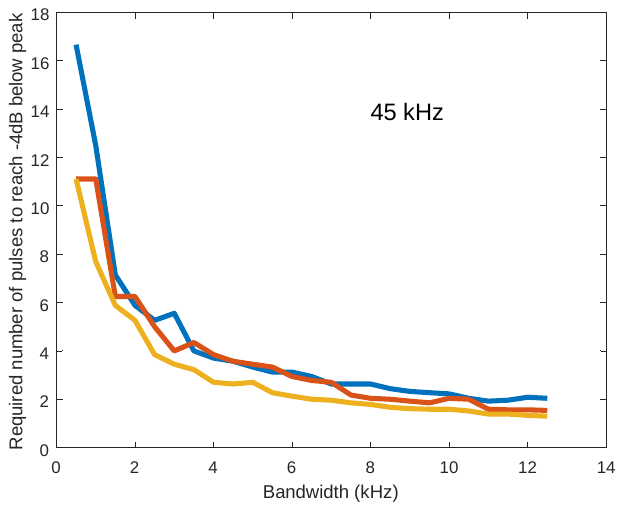

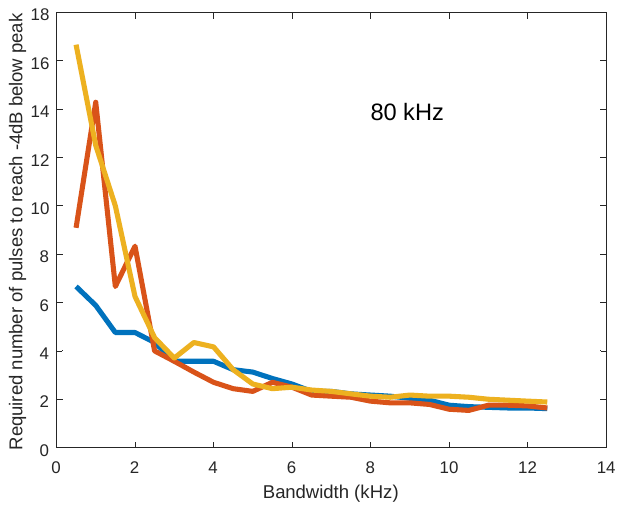

**Figure S9. Trading bandwidth for pulse repetition in order to increase detection sensitivity.** A bat can either increase its bandwidth (Figure 3 this paper) or it can repeat the emission several times to reach the same detection probability of detecting an insect swarm. Blue: swarm size N=16; Red: N=50; Orange: N=200. All graphs representing R=35mm. Upper frequencies of a signal with 12.5 kHz bandwidth are indicated within each graph. This graph shows how the two entities (bandwidth and repetition) relate to each other. When using bandwidths under 1.5 kHz, the required number of emissions to detect a swarm (detection level defined as -4dB below the peak level the swarm can evoke) quickly rises to above 10. In practice this means that the bat is losing time and energy (in having to emit more pulses) when not widening bandwidth.
